# Maternal immune activation and adolescent alcohol exposure increases alcohol drinking and disrupts cortical-striatal-hippocampal oscillations in adult offspring

**DOI:** 10.1101/2022.03.03.482905

**Authors:** Angela M Henricks, Emily DK Sullivan, Lucas L Dwiel, Judy Y Li, Diana J Wallin, Jibran Y. Khokhar, Wilder T Doucette

## Abstract

Maternal immune activation (MIA) is strongly associated with an increased risk of developing mental illness in adulthood, which often co-occurs with alcohol misuse. The current study aimed to begin to determine whether MIA, combined with adolescent alcohol exposure (AE), could be used as a model with which we could study the neurobiological mechanisms behind such co-occurring disorders. Pregnant Sprague-Dawley rats were treated with PolyI:C or saline on gestational day 15. Half of the offspring were given continuous access to alcohol during adolescence, leading to four experimental groups: controls, MIA, AE, and Dual (MIA + AE). We then evaluated whether MIA and/or AE alters: 1) alcohol consumption; and 2) cortical-striatal-hippocampal oscillations in adult offspring. Dual rats, particularly females, drank significantly more alcohol in adulthood compared to all other groups. Using machine learning to build predictive models from oscillations, we were able to differentiate Dual rats from control rats and AE rats in both sexes, and Dual rats from MIA rats in females. The current data suggest that MIA+AE (Dual “hits”) is a valuable model that we can use to study the neurobiological underpinnings of co-occurring disorders. Our future work aims to extend these findings to other addictive substances to enhance the translational relevance of this model.

## Introduction

Approximately 9.5 million adults in the United States are living with both a substance use disorder (SUD) and mental illness [1], typically referred to as “co-occurring disorders.” Alcohol is one of the most commonly abused substances in this population, with approximately 32% of individuals with mental illness misusing alcohol [1]. Individuals with co-occurring disorders are less responsive to treatment and experience higher rates of relapse, homelessness, incarceration, and suicide compared to individuals with a single disorder [2]. One reason for this difference in morbidity is because co-occurring disorders are notoriously difficult to treat and require integrative care, and the available pharmacotherapies are largely ineffective [3,4]. A major barrier to the development of better therapies is that there is still much to learn regarding the neurobiological mechanisms underlying these co-occurring disorders [5,6].

From a neurobiological perspective, it has been consistently demonstrated that the brain is highly susceptible to the harmful effects of environmental stressors in the early stages of development. One of these environmental risk factors is prenatal exposure to infection [7,8,9,10]. Systemic viral infections, like influenza or rubella, in pregnant women have been repeatedly associated with an increased incidence of psychosis- and mood-related disorders in offspring (e.g., schizophrenia, bipolar disorder, and depression) [8,9], and these mental illnesses are often comorbid with alcohol misuse [1]. However, since not all individuals exposed to infection in the prenatal environment develop a mental illness, it is likely that prenatal stressors combined with a “second-hit” during other critical periods of development (e.g., adolescence) further increase the probability of developing a mental illness in adulthood [8,9,10]. There is evidence for this “two-hit” model, reviewed elsewhere [9,10], indicating that a possible adolescent stressor is alcohol and/or drug use. Using maternal immune activation (MIA) to mimic prenatal exposure to infection in rodents, we therefore tested the hypothesis that MIA combined with adolescent alcohol exposure (AE) might serve as a useful heuristic with which we can begin to study co-occurring disorders.

We further hypothesized that MIA and/or AE would disrupt neural circuit activity in regions that regulate reward-related behaviors. Our previous clinical work has demonstrated that individuals with schizophrenia and co-occurring substance use disorder have reduced functional connectivity between the nucleus accumbens (NAc) and medial prefrontal cortex (mPFC) [11], which could serve as a therapeutic target in future research [12]. Another highly relevant region is the dorsal hippocampus, which plays a vital role in regulating emotions and facilitating learning, including responding to drug cues [13]. Further, MIA offspring have shown reduced connectivity between the mPFC and the CA1 of the hippocampus that correlates with abnormal prepulse inhibition, a well-characterized symptom of MIA exposure consistent with sensorimotor gating deficits observed across multiple psychiatric illnesses [14].

The current set of experiments thus investigated the impact of MIA, AE, and MIA+AE (Dual) on alcohol drinking behavior and local field potentials (LFPs) recorded from the mPFC, NAc shell, and CA1 in male and female rats. We aimed to determine: 1) whether MIA and/or AE alters alcohol consumption; 2) whether cortical-striatal-hippocampal oscillations could predict MIA and/or AE exposure.

## Materials and Methods

### General Experimental Design

<u>Experiment 1</u> allowed us to evaluate the impact of MIA and AE on adulthood alcohol drinking behavior. Male and female offspring were divided into four groups: control, MIA, AE, or Dual. Since MIA rats have shown both hypo- and hyperactive locomotor behavior in previous studies [15,16,17], we also tested rats’ locomotor response to a novel environment in adulthood to verify that our procedures caused a known behavioral phenotype of MIA. Rats were then trained to drink alcohol in their home cage, as described below.

<u>Experiment 2</u> allowed us to evaluate the impact of MIA and AE on cortical-striatal-hippocampal oscillations. A separate group of adult male and female control, MIA, AE, or Dual rats were implanted with electrodes targeting the prelimbic (PL) and infralimbic (IL) mPFC, NAc shell, and CA1. Following recovery, each rat underwent two, 30-minute recording sessions to measure baseline neural circuit activity, and we used an unbiased machine-learning approach to determine whether on cortical-striatal-hippocampal oscillations could predict MIA and/or AE exposure. See Figure 1 for the experimental timelines.

**Figure 1.**
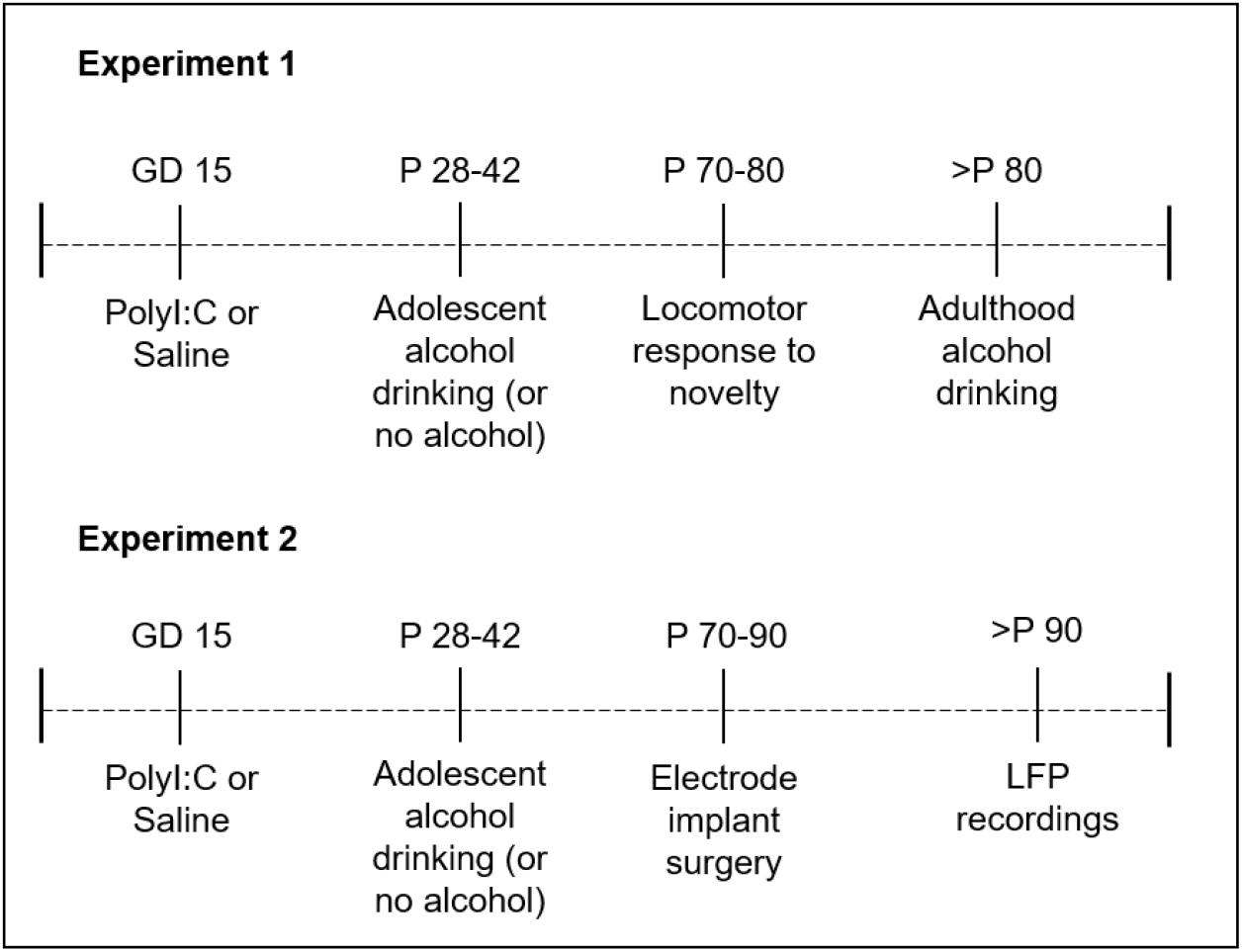
Experimental timelines.

### Animals

Timed-pregnant Sprague-Dawley rats were ordered to arrive on gestational day (GD) 8 (Charles River), and allowed to acclimate to the housing environment for 7 days prior to experimentation. Dams were housed individually on a reverse 12-hour light cycle with *ad libitum* access to food and water. Weaned pups were housed in same-sex pairs until postnatal day (P) 28, after which all animals were individually housed. All experiments were carried out in accordance with the National Institute of Health *Guide for the Care and Use of Laboratory Animals* (NIH Publications No. 80-23) and were approved by the Institutional Animal Care and Use Committee of Dartmouth College.

### Maternal Immune Activation

Pregnant dams were injected polyinosinic:polycytidylic acid [PolyI:C, 4mg/kg, IV] (Tocris Bioscience) or saline (1 mL/kg, IV) on GD 15. PolyI:C is a synthetic analog of double-stranded RNA, which leads to a heightened immune response in rats [10]. Dams’ body weight, food and water intake were monitored at -24, 0, 24, and 48 hours from injection. Blood was collected 2 hours after injection to measure the pro-inflammatory cytokines IL6 and TNFα. Both male and female pups were allowed to develop normally and weaned on P 21.

### Cytokine ELISA

Serum samples from dams were analyzed via enzyme linked immunosorbent assay (ELISA) for IL6 and TNFα (Thermo Fisher Scientific) following manufacturer’s recommendations.

### Adolescent Alcohol Exposure

AE and Dual rats were allowed to drink 10% alcohol in their home cage from P 28-42, consistent with our previous work [18]. Rats were given 24 hours access to alcohol and water, and the weight of each bottle was measured daily. The position of each bottle was rotated daily to avoid positional preference, and rats were weighed weekly to calculate weight-adjusted alcohol consumption (g/kg).

### Locomotor Response to a Novel Environment

On approximately P 70, rats were allowed to explore a novel open field (60cm x 60cm x 33cm) in the dark for 25 minutes. Infrared cameras recorded the behavior, which was analyzed with EthoVision behavioral tracking software (Noldus Information Technology). Total distance traveled was calculated and used for data analysis.

### Adulthood Alcohol Drinking

On approximately P 80, rats were trained to drink 10% alcohol in their home cage for 90 min/day, 5 days/week using a sucrose fade technique like that described previously [19]. Briefly, rats were allowed to drink 5% sucrose in water during week 1, then 5% sucrose + 10% alcohol during week 2, then 2.5% sucrose + 10% alcohol during week 3, then only 10% alcohol in water for three more weeks. Rats were weighed weekly to assess amount of alcohol consumed (g/kg).

### Surgery

Electrodes were designed and constructed in-house and were similar to those used in our previous publications [20,21,22]. Animals were anesthetized with isoflurane gas (4% induction, 2% maintenance) and mounted in a stereotaxic frame. Custom electrodes were implanted bilaterally targeting the PL mPFC (from bregma: DV -4mm; AP +3.4mm; ML +/-0.75mm), IL mPFC (from bregma: DV -5mm; AP +3.4mm; ML +/-0.75mm), NAc shell (from bregma: DV - 8mm; AP +1.2mm; ML +/-1.0mm), and CA1 of the hippocampus (from bregma: DV -2.5mm; AP -3.8mm; ML +/-2.5mm). Four stainless steel skull screws were placed around the electrode site and dental cement (Dentsply) was applied to secure the electrodes in place. Rats were allowed to recover for at least 7 days before any experimentation began.

#### Histology

At the end of the experiments, rats were euthanized via CO_2_ gas inhalation. Brains were harvested from rats implanted with electrodes and flash frozen in 2-methylbutane on dry ice. Tissue was stored at -20 °C prior to being sectioned at 50 μm using a Leica CM1850 cryostat and stained with thionin. Electrode placement was verified using a Leica A60 microscope. Out of the 84 rats implanted, we were unable to check histology on a small number of brains due to tissue damage that occurred during the collection process: 3 whole brains (2 control and 1 Dual); mPFC and/or NAc regions from 5 brains (1 control, 1 MIA, 2 AE, and 1 Dual). However, since no other brains were removed due to misplaced electrodes, we used all data for analysis. Figure 5A depicts representations of electrode placements.

### Local Field Potential Recordings

LFPs were recorded from each awake, freely-behaving rat in a standard operant chamber (MedAssociates). Rats were allowed to move about the chamber, but there was no task involved and rats did not have any prior experience in the chambers. Data from each recording were analyzed using frequency ranges from the rodent literature (δ =1-4 Hz, θ = 5-10 Hz, α = 11-14 Hz, β = 15-30 Hz, low γ = 45-65 Hz, and high γ = 70-90 Hz). LFP signal processing to characterize the power spectral densities (PSD) within, and coherence between brain regions, for each rat was calculated using custom code written for Matlab, as we have previously published [20,21,22]

### Statistical Analysis

#### Dam analysis

Pro-inflammatory cytokine levels were analyzed using an independent samples t-tests comparing Control and MIA dams. A repeated measures ANOVA was used to compare behavioral and physiological responses to PolyI:C using time as the within-subject variable and group (Control or MIA) as the between-subject variable.

#### Behavioral analysis in offspring

The average total distance traveled was calculated for the entire locomotor response to novelty session. A three-way ANOVA analyzed the impact of MIA, AE, and sex. The average g/kg of alcohol consumed in adolescence and adulthood was analyzed using a repeated measures ANOVA, with time as the within-subject variable and MIA, AE, and sex as the between-subject variables.

#### LFP oscillation analysis

Similar to our previous publications [20,21,22] we built general linear models to classify rats based on group assignment (i.e., control, MIA, AE, or Dual) using LFP oscillation data. Since each rat underwent two baseline recording sessions, we used data from both sessions to build baseline models. Data were then calculated in 5 second bins, with each bin representing one sample in the models. Using a “leave-one-out” (LOO) approach, models were then trained on all data minus one animal from each group, and then the model was tested on the left-out animal. To account for overrepresenting animals with more “clean” data (i.e., low noise) than other animals with less “clean” data (i.e., high noise), each model used only 1200 samples from each animal. All possible combinations of LOO were analyzed, and each LOO combination was run 100 times to account for sub-sampling. Model performance is reported as mean area under the receiver operating characteristic curve (AUC) ± 95% confidence interval.

Because we used multiple recording sessions from the same animal and 5-second bins as samples, we also evaluated models built on permutations of binary rat groupings (“animal detector”), as previously described [22]. This was done by keeping the LFP oscillation data together with the rat it was recorded from, and shuffling the group assignment of each rat’s set of recordings. The “animal detector” test thus allowed us to determine how much of our model accuracy was simply due to the ability of the algorithm to predict individual differences in oscillations *not* related to group assignment. We then compared the model performance of the real data to the “animal detector.” If the real model performed with greater accuracy than the “animal detector,” it indicated that information existed in the LFP signal. We then implemented exhaustive single feature regressions using each LFP predictor to determine the relative information content of each neural feature. Figure 5B depicts the model building approach.

## Results

### The Impact of PolyI:C on Dams

PolyI:C did not influence the total number of days in gestation, the number of pups born, or the M/F pup ratio (all *p* values > 0.05; Figure 2A and 2B). However, polyI:C did reduce both food and water intake, and slowed weight gain for 24 hours compared to saline (all *p* values < 0.05; Figure 2C, 2D, and 2E). While polyI:C did not significantly enhance circulating IL6 or TNFα two hours after injection, there was a trend for IL6 levels to be increased in PolyI:C dams [*t*(7) = - 2.04, *p* = 0.08; Figure 2F]. Nevertheless, the reduced weight gain and eating behavior of the dams exposed to PolyI:C is consistent with other reports [16,17], and combined with the hypolocomotion we observed in MIA offspring (see below), we are confident that PolyI:C induced MIA in our study. In future studies, collecting blood at different timepoints (e.g., 3-6 hours) may help capture the previously observed increases in pro-inflammatory cytokines [23].

**Figure 2.**
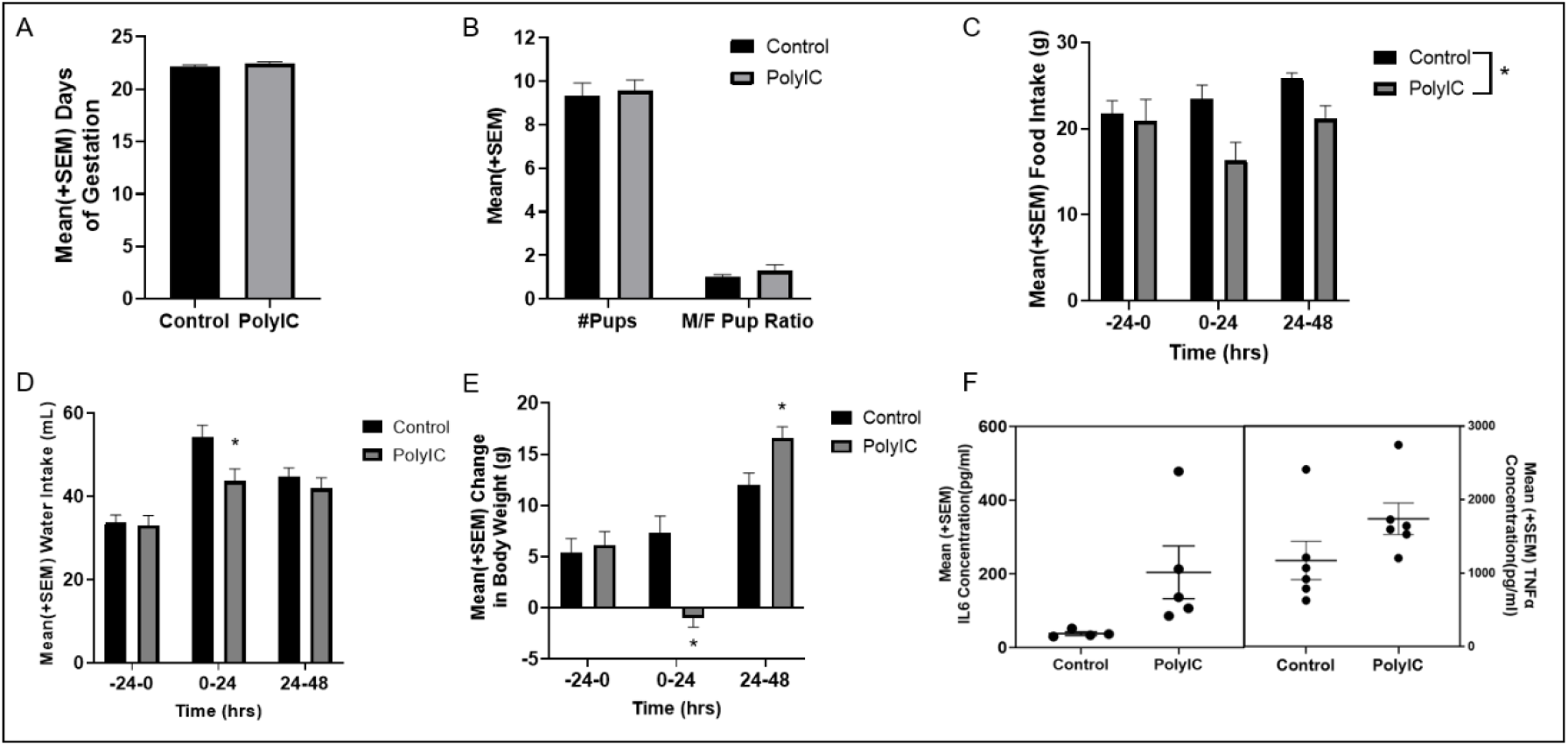
The impact if PolyI:C on dams. **A**) PolyI:C did not impact the number of days of gestation compare to saline-treated dams (Controls) (p > 0.05). **B**) PolyI:C did not impact the number of pups born or M/F pup ratio compare to Controls dams (p > 0.05). **C**) Dams treated with PolyI:C ate less food than Control dams overall (p < 0.05). **D**) Dams treated with PolyI:C drank less water than Control dams 24 hours after injection (p < 0.05). **E**) PolyI:C caused a reduction in body weight 24 hours after injection, but an increase 48 hours after injection, compared to Control dams (p < 0.05). **F**) The impact of PolyI:C on IL6 and TNFα concentration 2 hours post-injection. Each dot represents an individual dam.

### Experiment 1: The impact of MIA and AE on behavior

#### Alcohol Consumption in Adolescence

A repeated measures ANOVA revealed a significant effect of time [*F*(6.01, 210.31) = 5.80, *p* < 0.01, *n*^*2*^*p* = 0.14; Figure 3A], but no effect of MIA, sex, an MIA*sex interaction, or a time*sex*MIA interaction (all *p* values > 0.05; n = 9-11/group/sex). While there was a significant time*sex interaction [*F*(6.01, 210.31) = 2.70, *p* < 0.05, *n*^*2*^*p* = 0.07], *post-hoc* analyses did not reveal any sessions in which there was a significant difference in alcohol consumed between male and female rats (all *p* values > 0.05; Figure 3A).

**Figure 3.**
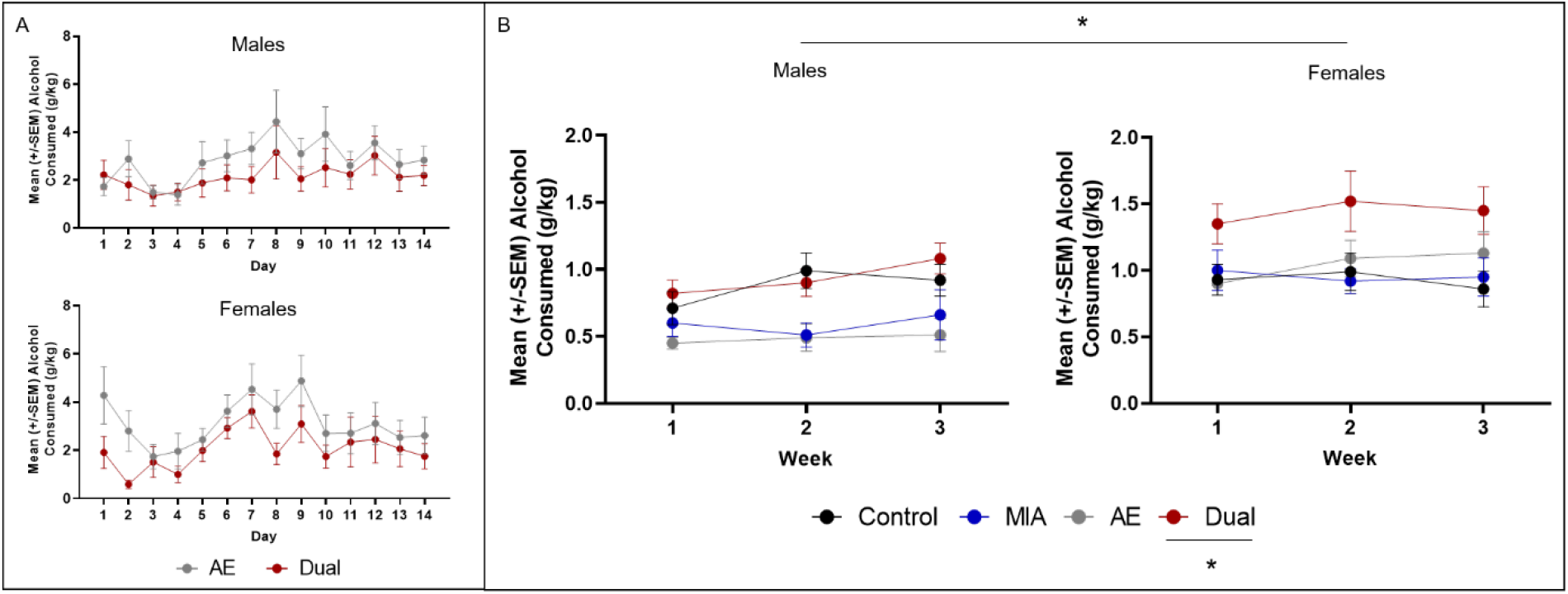
The impact of MIA and AE on offspring drinking behavior. **A**) Average alcohol consumed (g/kg) during adolescence (P 28-42) in male and female offspring from dams exposed to PolyI:C (Dual) or saline (AE). There were no significant differences between groups (*p* > 0.05). **B**) Average alcohol consumed (g/kg) in a limited access paradigm across 3 weeks in adult control, MIA, AE, and Dual rats. Drinking rates increased across time in all groups, but Dual offspring drank significantly more alcohol than all other groups (*p* < 0.05). Female rats drank more alcohol than males (*p* < 0.05).

#### Locomotor Response to Novelty in Adulthood

A three-way ANOVA revealed a significant effect of MIA [*F*(1, 73) = 10.27, *p* < 0.01, *n*^*2*^*p* = 0.12], sex [*F*(1, 73) = 32.56, *p* < 0.01, *n*^*2*^*p* = 0.31], and a MIA*sex*AE interaction [*F*(1, 73) = 4.16, *p* < 0.05, *n*^*2*^*p* = 0.05; n = 9-11/group/sex). Further analyses revealed that male MIA and Dual rats moved less than Control and AE rats (*p* < 0.05), and female rats overall moved more than male rats (*p* < 0.05; Figure 4).

**Figure 4.**
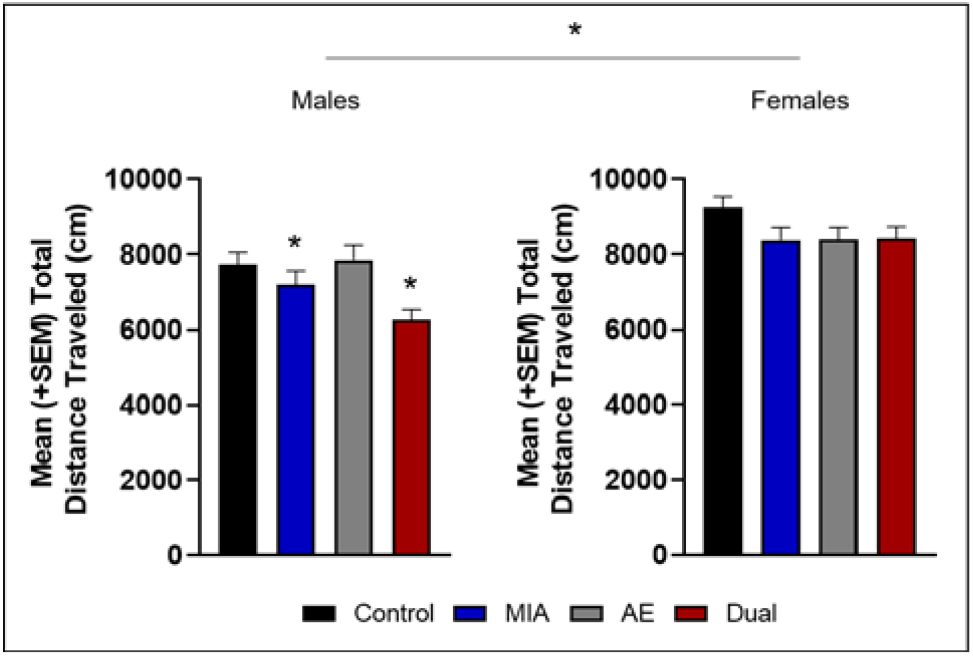
The impact of MIA and AE on offspring locomotor behavior. MIA and MIA+AE (Dual) reduced average distance traveled in a locomotor response to novelty task in male rats (*p* < 0.05). Female rats overall moved more than male rats (*p* < 0.05).

#### Alcohol Consumption in Adulthood

A repeated measures ANOVA revealed a significant effect of time [*F*(2, 136) = 4.41, *p* < 0.05, *n*^*2*^*p* = 0.06], a significant effect of sex [*F*(1, 68) = 19.11, *p* < 0.01, *n*^*2*^*p* = 0.22], and a significant MIA*AE interaction [*F*(1, 68) = 10.44, *p* < 0.01, *n*^*2*^*p* = 0.13; n = 8-11/group/sex]. Further analyses revealed that: 1) across weeks, alcohol consumption increased in all groups; 2) Dual rats drank more alcohol than Control, MIA, and AE rats, which was particularly apparent in females; and 3) female rats drank more alcohol than male rats (all *p* values < 0.05; Figure 3B). It is important to note that there was a trend for an AE*sex interaction [*F*(1, 68) = 3.62, *p* = 0.06, *n*^*2*^*p* = 0.05], and that while Dual male rats consumed more alcohol than the AE and MIA rats, they drank approximately the same amount as the control rats.

### Experiment 2: The impact of MIA and AE on LFP oscillations

In order to identify how MIA and AE impact cortical-striatal-hippocampal LFP oscillations, we built predictive models comparing Controls to MIA, AE, and Dual groups individually. LFP data were analyzed for each sex separately. Because the Dual rats showed significant increases in alcohol drinking compared to other groups, we used Dual rats as the comparison group. Using LFPs to predict Dual rats from Control rats, models for each sex outperformed the “animal detector” (Males real mean accuracy = 0.63±0.04; Females real mean accuracy: 0.68±0.05; Figure 5C). Further, models were able to predict Dual vs. MIA rats in females (real mean accuracy = 0.57±0.05), but not in males (real mean accuracy = 0.51±0.04; Figure 5D), and Dual vs. AE rats in both sexes (Males real mean accuracy = 0.55±0.06; Females real mean accuracy: 0.59±0.09; Figure 5E). The “animal detector” models estimated chance predictions in all cases, with a mean accuracy ranging from 0.47±0.04 – 0.52±0.04 (Figure 5C, 5D, and 5E).

**Figure 5.**
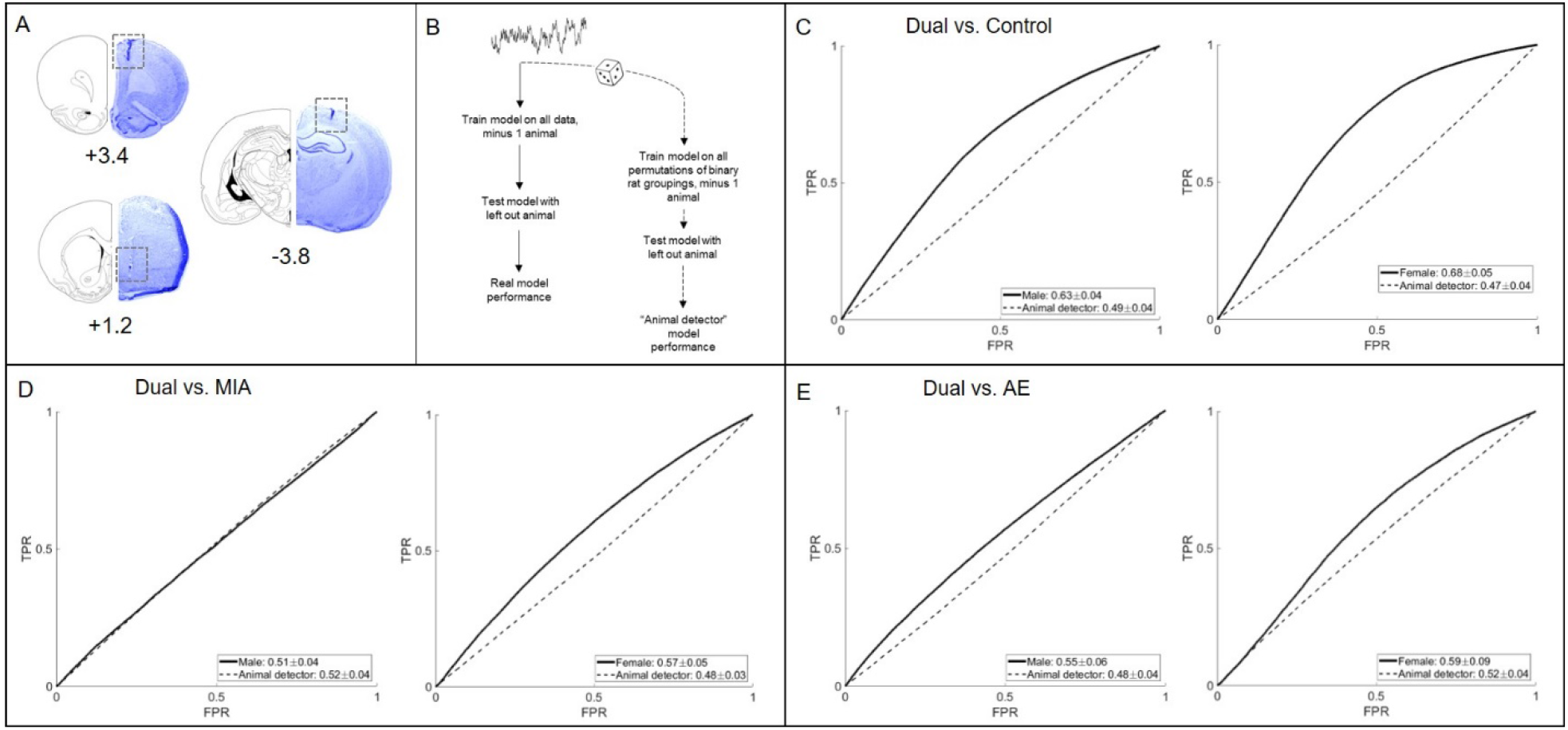
Cortical-striatal-hippocampal LFPs predict MIA and/or AE exposure. **A**) Histologic representation of lesions caused by electrode cannula in the mPFC (+3.4mm from bregma), NAcSh (+1.2mm from bregma), and CA1 (−3.8mm from bregma). Electrode wires extended 1mm from the end of the cannula (or 2mm for the IL mPFC). **B**) Schematic representation of the baseline model building. **C**) LFPs predicted Dual rats from control rats better than the “animal detector” in both males and females. **D**) LFPs predicted Dual rats from MIA rats better than the “animal detector” in females, but not in males. **E**) LFPs predicted Dual rats from AE rats better than the “animal detector” in both males and females.

Based on single-feature regression analyses, Table 1 shows the five features containing the most information (i.e., with the highest individual prediction accuracies) for each of the models that performed above chance estimates. It is interesting to note that all of the most predictive individual features are power features, and not coherence between regions.

**Table 1.** The 5 LFP features with the highest individual predictive accuracies of Dual vs. Control, Dual vs. MIA in females, and Dual vs. AE. Frequency bands [delta (δ), theta (θ), alpha (α), beta (β), low gamma (lγ), and high gamma (hγ)] are described for power features within neural sites. Left or right hemisphere is depicted as a subscript.

| Dual vs. Control |  | Dual vs. MIA |  | Dual vs. AE |  |
| --- | --- | --- | --- | --- | --- |
| Male | Female | Male | Female | Male | Female |
| PL<sub>R</sub> hγ | IL<sub>R</sub> lγ | ---- | IL<sub>R</sub> β | PL<sub>L</sub> δ | CA1<sub>L</sub> β |
| NAC<sub>R</sub> hγ | IL<sub>R</sub> δ | ---- | PL<sub>L</sub> β | CA1<sub>R</sub> β | NAC<sub>L</sub> β |
| IL<sub>L</sub> δ | IL<sub>R</sub> θ | ---- | IL<sub>R</sub> lγ | PL<sub>R</sub> δ | PL<sub>L</sub> β |
| IL<sub>L</sub> lγ | IL<sub>L</sub> lγ | ---- | IL<sub>R</sub> hγ | CA1<sub>L</sub> β | IL<sub>R</sub> β |
| CA1<sub>R</sub> δ | IL<sub>L</sub> δ | ---- | IL<sub>L</sub> β | PL<sub>R</sub> θ | IL<sub>R</sub> lγ |

## Discussion

MIA is a relatively well-characterized heuristic model in terms of the behavioral similarities to mental illnesses like schizophrenia, bipolar disorder, and depression [8]; MIA leads to maladaptive changes in locomotor behavior, affect, cognitive flexibility, sensorimotor gating, and social interactions [9,10]. However, even though mental illness often co-occurs with substance misuse, the current set of experiments are one of very few studies (discussed below) to test whether MIA rats might be more prone to addiction-like behaviors, and are thus a significant addition to the literature. Our data specifically indicate that MIA or AE alone does not impact alcohol drinking behavior, but that “two-hits” (e.g., MIA+AE; Dual) leads to enhanced alcohol consumption in adult offspring, particularly in females. Further, predictive models using cortical-striatal-hippocampal oscillations can differentiate Dual rats from controls and AE rats in both sexes, and Dual rats from MIA rats in females. These data suggest that there is information in these circuits regarding MIA and AE, and this information helps us begin to understanding the neurobiological underpinnings of the behavioral, cognitive, and sensorimotor deficits linked to prenatal exposure to infection and adolescent alcohol exposure [24,25].

While there are data to suggest that MIA alone alters reward behaviors in rodents, this work has almost entirely focused on motor responses to dopamine agonists [26,27,28]. Since the dopamine system is underactive in both schizophrenia and in MIA rats [29], it is unclear if these previous data represent an “addiction” phenotype, or are simply reflective of the locomotor deficits induced by the dopamine dysfunction. The present AE data also somewhat conflicts with previous reports, which show that AE increases alcohol consumption in adulthood [30,31; but see 32,33,34]. However, these studies offered alcohol intermittently, while the current study offered alcohol continuously in the home cage. We therefore hypothesize that AE in this study either primed MIA rats to drink alcohol in adulthood, helping eliminate the well-known taste aversion [35], or impacted neurodevelopment at a critical time-point that synergistically interacted with MIA to produce increased adulthood drinking (or both). As has been suggested previously [9,10], we hypothesize that “two-hits,” one in very early life and one in adolescence, are necessary to produce a phenotype that is similar to the clinical presentation of co-occurring disorders. The current data supports this hypothesis, and our future studies aim to advance this line of work by testing whether MIA and/or AE rats are willing to work harder for alcohol and other drugs than control rats in an operant setting.

It is important to highlight that we observed significant sex differences in these studies that may help us better understand the complex sex differences in clinical presentations of co-occurring disorders. Schizophrenia is more common in men, especially earlier in life, and men with schizophrenia are more likely to have a co-occurring SUD [36]. On the other hand, rates of alcohol misuse are increasing among women, and women experience more psychiatric issues related to alcohol misuse than men [37,38]. Our works aims to begin to disentangle the neurobiological contributions to these sex differences. To that aim, we replicated our previous studies showing that female rats drink more alcohol in general compared to males [19,22], and also add that females might be more susceptible the impact of the Dual hit on drinking behavior. Further, cortical-striatal-hippocampal oscillations were not predictive of Dual vs. MIA rats in males, as it was in females. These data suggest that the adolescent alcohol exposure in MIA males may dysregulate activity in circuits not measured in these studies. Other regions involved in decision making and stress, like the orbital-frontal cortex and amygdala, are impacted on both a structural and functional level by adolescent alcohol exposure [39], and thus might contain more information regarding Dual exposure in males.

Further, the primary neural features that differentiated across groups were sexually dimorphic. Predictive models distinguished Dual rats from control rats, and Dual rats from AE rats, relatively well in both sexes. However, an interesting pattern emerged when we looked at the most predictive neural features for each sex. Cortical features, particularly from the IL, largely contained the most information in predicting females from every other group. In males, the predictive features were more mixed; though a lot of the information still came from cortical sites, hippocampal and striatal features were also predictive of male Dual rats. These data help us identify which neural features to target to reduce alcohol misuse in future studies. For example, we have previously shown that deep brain stimulation to the NAc shell has the capacity to reduce alcohol consumption in high-drinking male rats [21], and these data are only a small portion of the literature suggesting the neuromodulation-based therapies can be efficacious for addiction and other neuropsychiatric disorders [40]. However, these studies have almost exclusively been done in male animals. In order to successfully initiate neuromodulation-based therapies that will be relevant to clinical populations, it is vital to understand “where” and “what” (e.g., striatal vs. cortical oscillations) we should be targeting, and these parameters may be different in males vs. females. The present study is a promising first step in the process of identifying neural features that may underly maladaptive behaviors, and our future work will test the capacity for cortical vs. striatal stimulation to reduce drinking in male and female Dual rats.

## Conclusion

The data presented herein provide support for a novel heuristic neurodevelopmental model which we can use to study the neurobiological basis of co-occurring mental illness and substance use. Although the current study is specifically focused on alcohol drinking, these results may also have significant implications for other addictive substances like cannabis and nicotine, which are two of the other most commonly used drugs in individuals with mental illness [1]. Thus, the Dual model in the present study may serve as a translationally-relevant platform on which we can better study the neurobiological underpinnings of, and develop treatments for, co-occurring disorders. These studies and our future work will contribute to the larger goal of identifying how early environmental stressors change the brain in such a way as to predispose an individual to develop a mental illness and/or SUD, uncovering neurobiological treatment targets for therapeutic development.

## Acknowledgments

The authors would like to acknowledge Dr. Alan I. Green, who passed away in November of 2020, and without whom these studies could not have been performed. Dr. Green was an incredible mentor to all of the authors, and he will be greatly missed for his brilliant insight, wit, and generosity.

We would also like to thank lab manager Elise Bragg for her assistance in collecting portions of the behavioral and histological data.

This work was supported by funds from the Department of Psychiatry at the Geisel School of Medicine at Dartmouth (AIG), the Hitchcock Foundation (AMH), NIDA T32 training grant (DA037202; AMH), an LRP grant from NIH NCATS (WTD), a NIAAA training grant (F31AA027441; LLD), and the Dartmouth Clinical and Translational Science Institute from NIH NCATS (KL2TR001088; WTD).

## Author Contributions

AMH, JYK, and WTD developed the conceptual framework, hypotheses, and design of these experiments. AMH, EDKS, LLD, DJW, and JYL collected and analyzed these data. AMH wrote the manuscript, and all other authors contributed to editing.

## References

1. Substance Abuse and Mental Health Services Administration. (2020). Key substance use and mental health indicators in the United States: Results from the 2019 National Survey on Drug Use and Health (HHS Publication No. PEP20-07-01-001, NSDUH Series H-55). Rockville, MD: Center for Behavioral Health Statistics and Quality, Substance Abuse and Mental Health Services Administration.

2. Hunt GE, Siegfried N, Morley K, Brooke-Sumner C, Cleary M. (2019). Psychosocial interventions for people with both severe mental illness and substance misuse. Cochrane Database Syst Rev. 12(12):CD001088.

3. Flynn PM, Brown BS. (2008). Co-occurring disorders in substance abuse treatment: issues and prospects. J Subst Abuse Treat. 34(1):36–47.

4. Morojele NK, Saban A, Seedat S. (2012). Clinical presentations and diagnostic issues in dual diagnosis disorders. Curr Opin Psychiatry. 25(3):181–186.

5. Balhara YP, Kuppili PP, Gupta R. (2017). Neurobiology of Comorbid Substance Use Disorders and Psychiatric Disorders: Current State of Evidence. J Addict Nurs. 28(1):11–26.

6. Gómez-Coronado N, Sethi R, Bortolasci CC, Arancini L, Berk M, Dodd S. (2018). A review of the neurobiological underpinning of comorbid substance use and mood disorders. J Affect Disord. 241:388–401.

7. Miguel PM, Pereira LO, Silveira PP, Meaney MJ. (2019). Early environmental influences on the development of children’s brain structure and function. Dev Med Child Neurol. 61(10):1127–1133.

8. Brown AS, Meyer U. (2018). Maternal Immune Activation and Neuropsychiatric Illness: A Translational Research Perspective. Am J Psychiatry. 175(11):1073–1083.

9. Estes ML, McAllister AK. (2016). Maternal immune activation: Implications for neuropsychiatric disorders. Science. 353(6301):772–7.

10. Meyer U. (2014). Prenatal poly(i:C) exposure and other developmental immune activation models in rodent systems. Biol Psychiatry. 75(4):307–15.

11. Fischer AS, Whitfield-Gabrieli S, Roth RM, Brunette MF, Green AI. (2014). Impaired functional connectivity of brain reward circuitry in patients with schizophrenia and cannabis use disorder: Effects of cannabis and THC. Schizophr Res. 158(1-3):176-82.

12. Khokhar JY, Dwiel LL, Henricks AM, Doucette WT, Green AI. (2018). The link between schizophrenia and substance use disorder: A unifying hypothesis. Schizophr Res., 194:78–85.

13. Jasinska AJ, Stein EA, Kaiser J, Naumer MJ, Yalachkov Y. (2014). Factors modulating neural reactivity to drug cues in addiction: a survey of human neuroimaging studies. Neurosci Biobehav Rev., 38:1–16.

14. Dickerson DD, Wolff AR, Bilkey DK. (2010). Abnormal long-range neural synchrony in a maternal immune activation animal model of schizophrenia. J Neurosci. 30(37):12424–31.

15. Batinić B, Santrač A, Divović B, Timić T, Stanković T, Obradović ALj, Joksimović S, Savić MM. (2016). Lipopolysaccharide exposure during late embryogenesis results in diminished locomotor activity and amphetamine response in females and spatial cognition impairment in males in adult, but not adolescent rat offspring. Behav Brain Res., 15;299:72-80.

16. Howland JG, Cazakoff BN, Zhang Y. (2012). Altered object-in-place recognition memory, prepulse inhibition, and locomotor activity in the offspring of rats exposed to a viral mimetic during pregnancy. Neuroscience, 10;201:184–98.

17. Vorhees CV, Graham DL, Braun AA, Schaefer TL, Skelton MR, Richtand NM, Williams MT. (2015). Prenatal immune challenge in rats: effects of polyinosinic-polycytidylic acid on spatial learning, prepulse inhibition, conditioned fear, and responses to MK-801 and amphetamine. Neurotoxicol Teratol., 47:54–65.

18. Khokhar JY, Todd TP. (2018). Behavioral predictors of alcohol drinking in a neurodevelopmental rat model of schizophrenia and co-occurring alcohol use disorder. Schizophr Res., 194:91–97.

19. Henricks AM, Berger AL, Lugo JM, Baxter-Potter LN, Bieniasz KV, Craft RM, McLaughlin RJ. (2016). Sex differences in alcohol consumption and alterations in nucleus accumbens endocannabinoid mRNA in alcohol-dependent rats. Neuroscience, 335:195–206.

20. Doucette WT, Dwiel L, Boyce JE, Simon AA, Khokhar JY, Green AI. (2018). Machine Learning Based Classification of Deep Brain Stimulation Outcomes in a Rat Model of Binge Eating Using Ventral Striatal Oscillations. Front Psychiatry, 9:336.

21. Henricks AM, Dwiel LL, Deveau NH, Simon AA, Ruiz-Jaquez MJ, Green AI, Doucette WT. (2019). Corticostriatal Oscillations Predict High vs. Low Drinkers in a Rat Model of Limited Access Alcohol Consumption. Front Syst Neurosci., 13:35.

22. Henricks AM, Sullivan EDK, Dwiel LL, Keus KM, Adner ED, Green AI, Doucette WT. (2019). Sex differences in the ability of corticostriatal oscillations to predict rodent alcohol consumption. Biol Sex Differ., 10(1):61.

23. Cunningham C, Campion S, Teeling J, Felton L, Perry VH. (2007). The sickness behaviour and CNS inflammatory mediator profile induced by systemic challenge of mice with synthetic double-stranded RNA (poly I:C). Brain Behav Immun. 21(4):490–502.

24. Gumusoglu SB, Stevens HE. (2019). Maternal Inflammation and Neurodevelopmental Programming: A Review of Preclinical Outcomes and Implications for Translational Psychiatry. Biol Psychiatry, 85(2):107–121.

25. Spear LP. Effects of adolescent alcohol consumption on the brain and behaviour. (2018). Nat Rev Neurosci., 19(4):197–214.

26. Zager A, Mennecier G, Palermo-Neto J. (2012). Maternal immune activation in late gestation enhances locomotor response to acute but not chronic amphetamine treatment in male mice offspring: role of the D1 receptor. Behav Brain Res., 232(1):30–6.

27. Borçoi AR, Patti CL, Zanin KA, Hollais AW, Santos-Baldaia R, Ceccon LM, Berro LF, Wuo-Silva R, Grapiglia SB, Ribeiro LT, Lopes-Silva LB, Frussa-Filho R. (2015). Effects of prenatal immune activation on amphetamine-induced addictive behaviors: Contributions from animal models. Prog Neuropsychopharmacol Biol Psychiatry. 63:63–9.

28. Straley ME, Van Oeffelen W, Theze S, Sullivan AM, O’Mahony SM, Cryan JF, O’Keeffe GW. (2017). Distinct alterations in motor & reward seeking behavior are dependent on the gestational age of exposure to LPS-induced maternal immune activation. Brain Behav Immun., 63:21–34.

29. Luchicchi A, Lecca S, Melis M, De Felice M, Cadeddu F, Frau R, Muntoni AL, Fadda P, Devoto P, Pistis M. (2016). Maternal Immune Activation Disrupts Dopamine System in the Offspring. Int J Neuropsychopharmacol. 19(7):pyw007.

30. Gass JT, Glen WB Jr, McGonigal JT, Trantham-Davidson H, Lopez MF, Randall PK, Yaxley R, Floresco SB, Chandler LJ. (2014). Adolescent alcohol exposure reduces behavioral flexibility, promotes disinhibition, and increases resistance to extinction of ethanol self-administration in adulthood. Neuropsychopharmacology, 39(11):2570–83.

31. Pascual M, Boix J, Felipo V, Guerri C. (2009). Repeated alcohol administration during adolescence causes changes in the mesolimbic dopaminergic and glutamatergic systems and promotes alcohol intake in the adult rat. J Neurochem., 108(4):920–31.

32. Slawecki CJ, Betancourt M. (2002). Effects of adolescent ethanol exposure on ethanol consumption in adult rats. Alcohol, 26(1):23–30.

33. Hamidullah S, Lutelmowski CD, Creighton SD, Luciani KR, Frie JA, Winters BD, Khokhar JY. (2021). Effects of vapourized THC and voluntary alcohol drinking during adolescence on cognition, reward, and anxiety-like behaviours in rats. Prog Neuropsychopharmacol Biol Psychiatry, 106:110141.

34. Vetter CS, Doremus-Fitzwater TL, Spear LP. (2007). Time course of elevated ethanol intake in adolescent relative to adult rats under continuous, voluntary-access conditions. Alcohol Clin Exp Res., 31(7):1159–68.

35. Saalfield J, Spear L. (2015). Consequences of repeated ethanol exposure during early or late adolescence on conditioned taste aversions in rats. Dev Cogn Neurosci., 16:174–182.

36. Abel KM, Drake R, Goldstein JM. Sex differences in schizophrenia. (2010). Int Rev Psychiatry, 22(5):417–28.

37. Grant BF, Chou SP, Saha TD, Pickering RP, Kerridge BT, Ruan WJ, Huang B, Jung J, Zhang H, Fan A, Hasin DS. (2017). Prevalence of 12-Month Alcohol Use, High-Risk Drinking, and DSM-IV Alcohol Use Disorder in the United States, 2001-2002 to 2012-2013: Results From the National Epidemiologic Survey on Alcohol and Related Conditions. JAMA Psychiatry, 74(9):911–923.

38. Erol A, Karpyak VM. (2015). Sex and gender-related differences in alcohol use and its consequences: Contemporary knowledge and future research considerations. Drug Alcohol Depend, 1;156:1–13.

39. Crews FT, Vetreno RP, Broadwater MA, Robinson DL. (2016). Adolescent Alcohol Exposure Persistently Impacts Adult Neurobiology and Behavior. Pharmacol Rev., 68(4):1074–1109.

40. Sullivan CRP, Olsen S, Widge AS. (2021). Deep brain stimulation for psychiatric disorders: From focal brain targets to cognitive networks. Neuroimage. 225:117515.

